# Characterizing the Spatiotemporal Boundaries of Functional Ultrasound for Brain Decoding

**DOI:** 10.64898/2026.08.30.748086

**Authors:** Cheng Bian, Jin Yang, Muyun Xie, Jiaru He, Yin Hing Sophia Sun, Jianjian Zhao, Zihao Chen, Hsin-Yi Lai, Zhen Yuan, Yimin Wang, Zhihai Qiu

## Abstract

Functional ultrasound imaging (fUS) measures task-associated changes in cerebral blood volume (CBV) with fine spatial sampling and rapid acquisition, supporting functional mapping and brain-signal decoding. As an indirect measure of neural activity, fUS-CBV signals are shaped by neurovascular coupling and vascular dynamics, which influence the spatial and temporal stimulus information available for decoding. Consequently, the functional distinctions accessible in fUS-CBV signals require characterization under controlled experimental conditions. Here, controlled whisker stimulation in awake, head-fixed mice was used to characterize these decoding-relevant response properties. Whisker identity and number, stimulation duration, and inter-stimulus interval were systematically varied. All-whisker stimulation produced contralaterally dominant responses in the primary somatosensory barrel field and ventral posteromedial thalamic nucleus. Different whisker inputs were associated with distinguishable cortical response distributions, while stimulation involving more whiskers produced broader representative activation patterns. Across durations of 0.5–10 s, brief stimulation generated measurable stimulus-associated signals, and response magnitude, persistence, and spatial distribution varied with stimulus duration. For paired 2-s stimuli, trial-level separability was already high at the shortest tested 1-s gap, corresponding to a 3-s onset-to-onset spacing, and reached 100% at gaps of 3 s or longer. Building on the experimentally characterized spatiotemporal response properties of fUS-CBV signals, a proof-of-concept modeling and decoding framework was developed, comprising cross-animal spatial prediction, classification of short versus long stimulation from S1BF ΔCBV time courses with known stimulus onset, and modeling of paired-event separability. Together, the results show that fUS-CBV signals retain structured spatial and temporal information about controlled sensory input after neurovascular transformation. These measured response properties characterize the spatiotemporal boundaries within which neural activity can be reliably distinguished using fUS-CBV signals, providing constraints for the development of fUS-based decoding models and the design of stimulation paradigms suitable for reliable fUS decoding.

## Introduction

Functional ultrasound imaging (fUS) combines ultrafast plane-wave acquisition with power Doppler processing to measure relative changes in cerebral blood volume (CBV) across cortical and subcortical structures ^1,2^. Rapid acquisition, fine spatial sampling, and spatiotemporal clutter filtering provide sensitivity to CBV changes in small cerebral vessels ^3^. Simultaneous electrophysiological, calcium imaging, and fUS measurements have identified systematic relationships between ultrasound-derived vascular signals and locally measured neuronal activity ^4–6^. These relationships arise through neurovascular coupling and the coordinated responses of neurons, glial cells and different vascular compartments ^7,8^. These measurement capabilities have supported functional mapping in behaving macaques, including single-trial detection of task-related activity ^9^, offline decoding of movement direction and effector ^10^, online closed-loop decoding of multiple movement directions ^11^, and identification of behavioral and self-motion states ^12,13^. In mice, fUS has also been applied to brain-wide sensory mapping and the analysis of signals associated with experimentally defined neural circuits ^14,15^. Functional measurements have further been extended to awake and freely moving rodents ^16,17^. Clinical applications of fUS include neonatal brain monitoring, intraoperative functional mapping and bedside detection of auditory responses in comatose patients ^18–20^.

Although previous studies have demonstrated that functional brain states and behavioral variables can be detected or decoded from fUS signals in both rodents and non-human primates, these task-specific demonstrations of feasibility do not establish the fundamental spatial and temporal boundaries within which neural information remains reliably distinguishable after its transformation into a hemodynamic signal. For functional decoding, this distinction is critical because the technical sampling characteristics of fUS do not necessarily define its effective spatial and temporal decoding capabilities. fUS does not measure neuronal firing directly, but instead detects cerebral hemodynamic responses generated through neurovascular coupling. Neural activity is therefore transformed into a vascular signal before being captured by fUS, and this transformation is constrained by the spatial distribution, density, and architecture of the local vasculature, as well as by the sensitivity and dynamics of neurovascular coupling ^4–8^.

Consequently, a critical question for the development of fUS as a decoding modality—and ultimately for its use in acoustic brain–computer interfaces—is at what spatial and temporal scales neural activity remains distinguishable in fUS-CBV signals after this neurovascular transformation. This includes determining how brief a sensory input can be while still producing a detectable hemodynamic response, whether inputs represented by spatially distinct neural populations retain separable vascular representations, and how closely spaced events can occur while remaining temporally distinguishable in the resulting fUS-CBV signals. These properties define the effective spatial and temporal boundaries of the information accessible to fUS, rather than simply the sampling capabilities of the imaging system. Quantifying these boundaries under controlled sensory stimulation is therefore essential for identifying suitable decoding scenarios, designing robust and decodable experimental paradigms, and establishing realistic spatial and temporal constraints for subsequent decoding models.

The mouse whisker–barrel system provides a controlled biological model for examining these questions. Individual facial whiskers are represented within an ordered array of cortical barrels, and tactile information reaches the barrel-field primary somatosensory cortex through brainstem and thalamic relays, including the ventral posteromedial nucleus ^21–24^. Whisker preference follows an organized spatial distribution at the population level, while local neuronal responses can extend across neighboring whisker representations ^7,25,26^. Whisker identity, the number of stimulated whiskers, stimulation duration, and temporal spacing can therefore be varied within a consistent experimental preparation. This organization enables defined changes in sensory input to be related to the spatial and temporal characteristics retained in fUS-CBV signals.

Previous fUS studies have mapped whisker-evoked cortical and thalamic activity and extended these measurements to awake, freely moving, and volumetric imaging settings ^1,14,15,27,28^. These studies establish the feasibility of detecting whisker-associated fUS signals, while the information retained across systematically varied spatial and temporal stimulation conditions remains to be characterized within a common experimental framework. The present study addresses three related questions: whether different whisker inputs retain distinguishable cortical response distributions, whether isolated brief stimuli and broad differences in stimulus duration remain detectable in fUS-CBV signals, and whether responses to successive stimuli can be identified as separate temporal components across different inter-stimulus intervals. Whisker identity and number, stimulation duration, and inter-stimulus interval were varied in awake, head-fixed mice to examine these properties. The resulting measurements were further incorporated into a proof-of-concept, experimentally informed framework linking selected stimulus factors with spatial activation patterns, broad duration classes, and paired-event separability. This approach characterizes decoding-relevant spatial and temporal information retained after neurovascular transformation and provides experimentally measured constraints for stimulus-related fUS decoding.

## Results

To determine which stimulus-related spatial and temporal features remained accessible in fUS-CBV signals, the analyses were organized as a sequential evaluation. Conventional all-whisker stimulation first established that the imaging system captured lateralized fUS-CBV responses in the S1BF and VPM. Whisker identity and number were then varied to examine spatial response patterns, whereas stimulus duration and inter-stimulus interval were varied to assess brief-event detectability and paired-event separability, respectively. These observations were subsequently integrated into a proof-of-concept, experimentally informed encoding and decoding framework.

### 1. Whisker stimulation evokes lateralized fUS-CBV responses in cortical and thalamic somatosensory regions

To determine whether the imaging configuration could capture whisker-evoked fUS-CBV responses, unilateral all-whisker stimulation was delivered using a block-design paradigm. Coronal fUS images were acquired using the synchronized experimental system shown in Fig. 1A. Each recording comprised five 20-s stimulation periods separated by 40-s intervals, with 60-s baseline periods before and after the stimulation sequence (Fig. 1B). Responses were evaluated in the primary somatosensory barrel field (S1BF) and the ventral posteromedial thalamic nucleus (VPM).

**Figure 1.**
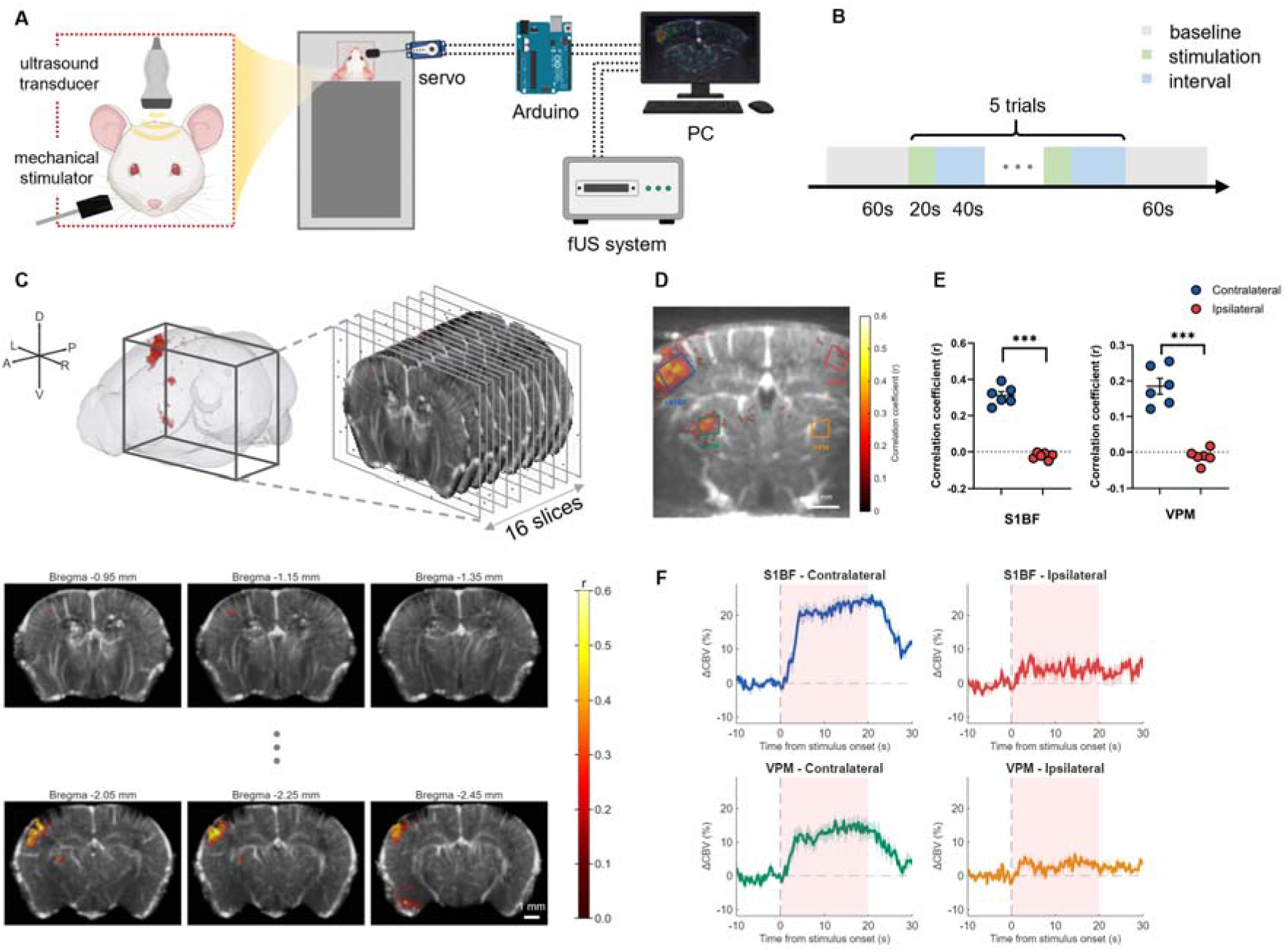
Experimental setup and validation of whisker-evoked fUS-CBV responses. **(A)** Schematic of the experimental setup for functional ultrasound imaging in awake, head-fixed mice during unilateral mechanical whisker stimulation. Whisker stimulation and fUS acquisition were synchronized using an Arduino-based control system. **(B)** Block-design stimulation paradigm. Each acquisition included an initial baseline period, five 20-s stimulation blocks separated by 40-s unstimulated intervals, and a final recovery period. **(C)** Atlas-space coverage of serial coronal fUS acquisitions. The three-dimensional view shows the spatial distribution and order of 16 acquired coronal planes. Six representative atlas-aligned correlation maps are shown below; the intervening planes are indicated by ellipses. **(D)** Representative atlas-aligned activation map with manually defined contralateral and ipsilateral regions of interest in the primary somatosensory barrel field (S1BF) and ventral posteromedial thalamic nucleus (VPM). Colors of ROI outlines correspond to the time courses in F. **(E)** Correlation coefficients within contralateral and ipsilateral S1BF and VPM ROIs across animals. Circles represent individual animals; horizontal bars indicate mean ± SEM. Asterisks indicate two-sided paired comparisons with Holm-adjusted P < 0.001. **(F)** Mean ΔCBV time courses in the four ROIs. Shaded areas indicate SEM, and pink shading indicates the stimulation period. Correlation maps in C and D are displayed using a common correlation-coefficient scale (n = 6 mice).

Voxel-wise correlation maps were generated by correlating the fUS-CBV time course with the corresponding stimulus regressor. In a representative experiment, positive correlations were localized predominantly to the contralateral somatosensory territory across multiple coronal sections. The atlas-aligned montage comprised 16 sections with atlas-derived Bregma coordinates ranging from −0.95 to −2.45 mm (Fig. 1C). A representative atlas-aligned coronal map showed stimulus-associated activation in the contralateral S1BF and VPM and defined bilateral S1BF and VPM regions of interest for subsequent analyses (Fig. 1D).

Group-averaged ΔCBV time courses showed stimulus-associated responses in the contralateral S1BF and VPM (Fig. 1F). In the contralateral S1BF, ΔCBV increased after stimulus onset, remained elevated during the 20-s stimulation period, and declined following stimulus cessation. The contralateral VPM showed a similar temporal profile with a smaller response amplitude. Responses in the ipsilateral S1BF and VPM remained comparatively weak over the same period.

Hemispheric differences were quantified using animal-level ROI correlation coefficients (Fig. 1E). For animals with multiple eligible coronal planes, correlation coefficients were Fisher-z transformed and averaged across planes before statistical testing. Across six animals, the correlation coefficient was higher in the contralateral S1BF than in the ipsilateral S1BF (0.311 ± 0.021 versus −0.023 ± 0.007, mean ± SEM; Holm-adjusted P = 1.11 × 10^−4^). A corresponding hemispheric difference was observed in the VPM (0.185 ± 0.022 versus −0.012 ± 0.008; Holm-adjusted P = 6.08 × 10^−4^).

These results show that unilateral whisker stimulation produced measurable and lateralized fUS-CBV responses in the S1BF and VPM. The contralaterally dominant cortical and thalamic responses established the experimental basis for examining how whisker identity, stimulation extent and temporal parameters were represented in subsequent fUS-CBV measurements.

### 2. Whisker identity and stimulation extent are reflected in spatial fUS-CBV response patterns

Following validation of lateralized responses in the S1BF and VPM, the spatial distributions associated with different whisker inputs were examined. Stimulation of a single C2 whisker produced localized positive correlations in the contralateral somatosensory cortex across consecutive coronal sections (Fig. 2A). Atlas-aligned three-dimensional reconstruction showed a spatially restricted cortical response in sagittal/oblique, coronal, and dorsal views. These results show that stimulation of an individual whisker produced a localized fUS-CBV response that remained identifiable across adjacent imaging planes.

**Figure 2.**
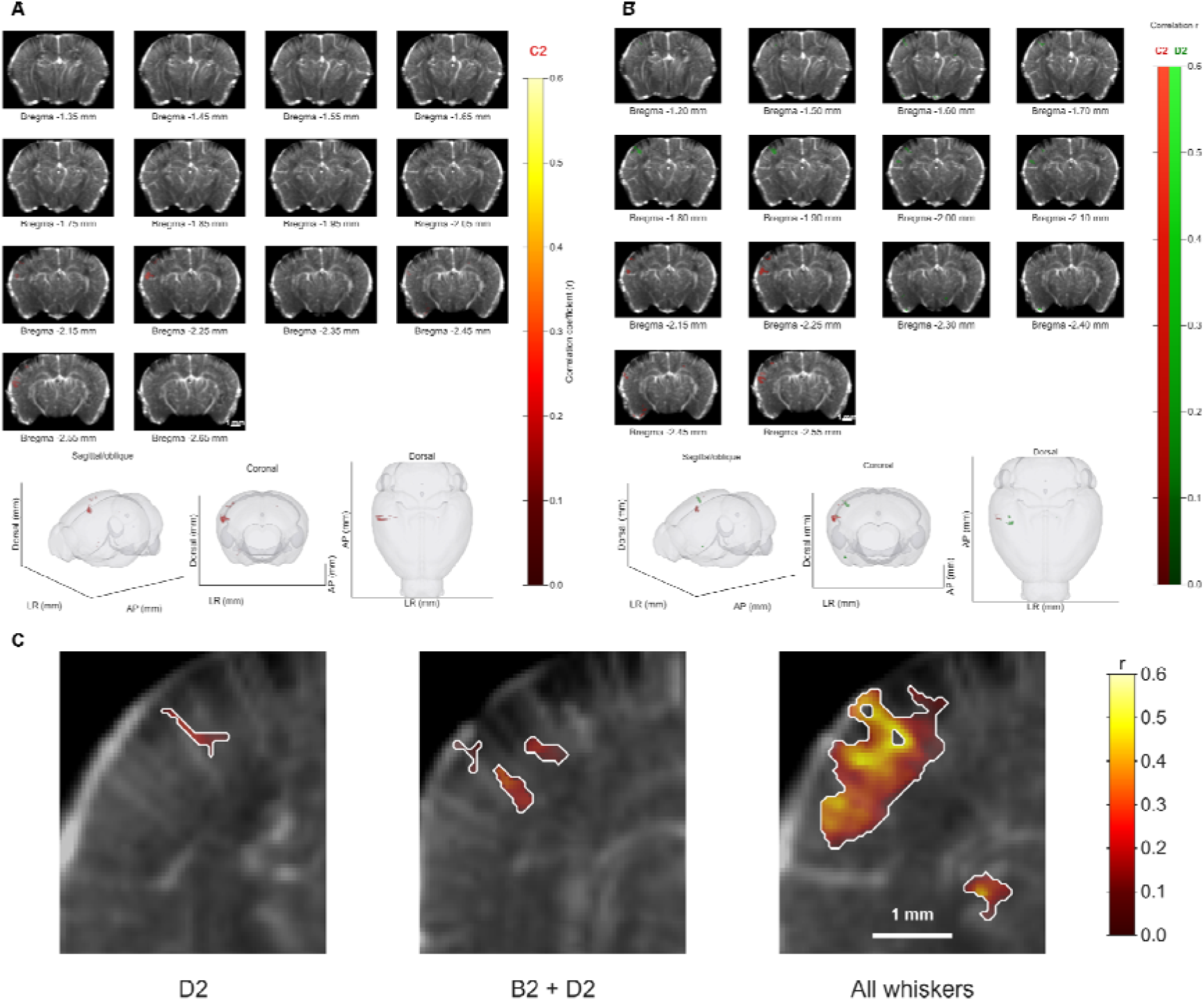
Spatial fUS-CBV response patterns associated with whisker identity and stimulation extent. **(A)** Representative C2-evoked activation maps displayed across serial coronal sections and in three-dimensional atlas-space views. Activation strength is represented by the correlation coefficient between the fUS-CBV signal and the stimulation regressor. **(B)** Dual-color overlay of D2- and C2-evoked activation maps, showing neighboring but spatially distinguishable cortical response distributions. Red denotes C2 and green denotes D2. **(C)** Representative atlas-aligned activation maps during single B2, combined B2+D2, and intact all-whisker stimulation. The maps are displayed at a matched coronal level using a common correlation scale and activation threshold, showing a progressive increase in activated cortical territory with the number of stimulated whiskers. Scale bars, 1 mm.

The spatial responses evoked by D2 and C2 stimulation were directly compared after alignment to the same atlas space. The two activation fields occupied neighboring but spatially distinct locations within the contralateral somatosensory cortex (Fig. 2B). Regions preferentially associated with either D2 or C2 stimulation remained visible in the dual-color overlay. These spatial differences show that fUS-CBV maps retained spatial information associated with whisker identity, with D2 and C2 stimulation producing distinguishable cortical response distributions.

The spatial extent of activation was subsequently compared across different whisker-stimulation conditions. Representative atlas-aligned maps from single B2 stimulation, combined B2+D2 stimulation, and stimulation of the intact whisker array were displayed at selected coronal planes using the same correlation scale and activation criterion (Fig. 2C). The activated cortical territory increased progressively from single-whisker stimulation to combined B2+D2 stimulation and stimulation of the intact whisker array.

Together, these results show that stimulation of different individual whiskers was associated with distinguishable cortical fUS-CBV response distributions. Conditions involving more stimulated whiskers showed broader response patterns in the representative maps. The measured spatial distributions therefore retained information related to both whisker identity and the extent of whisker stimulation.

### 3. Duration-related characteristics of whisker-evoked fUS-CBV responses

A stimulation-duration series ranging from 0.5 to 10 s was used to examine how the measured fUS-CBV response varied across whisker-stimulation durations (Fig. 3A). Stimulus-aligned S1BF ΔCBV traces showed measurable signal increases across the tested conditions (Fig. 3B). A response was already evident in the group-averaged trace following 0.5-s stimulation. Longer stimulation was generally accompanied by a greater ΔCBV increase and a more persistent elevation of the signal. Across durations, the fUS-CBV response developed after stimulus onset and extended beyond the corresponding mechanical stimulation period.

**Figure 3.**
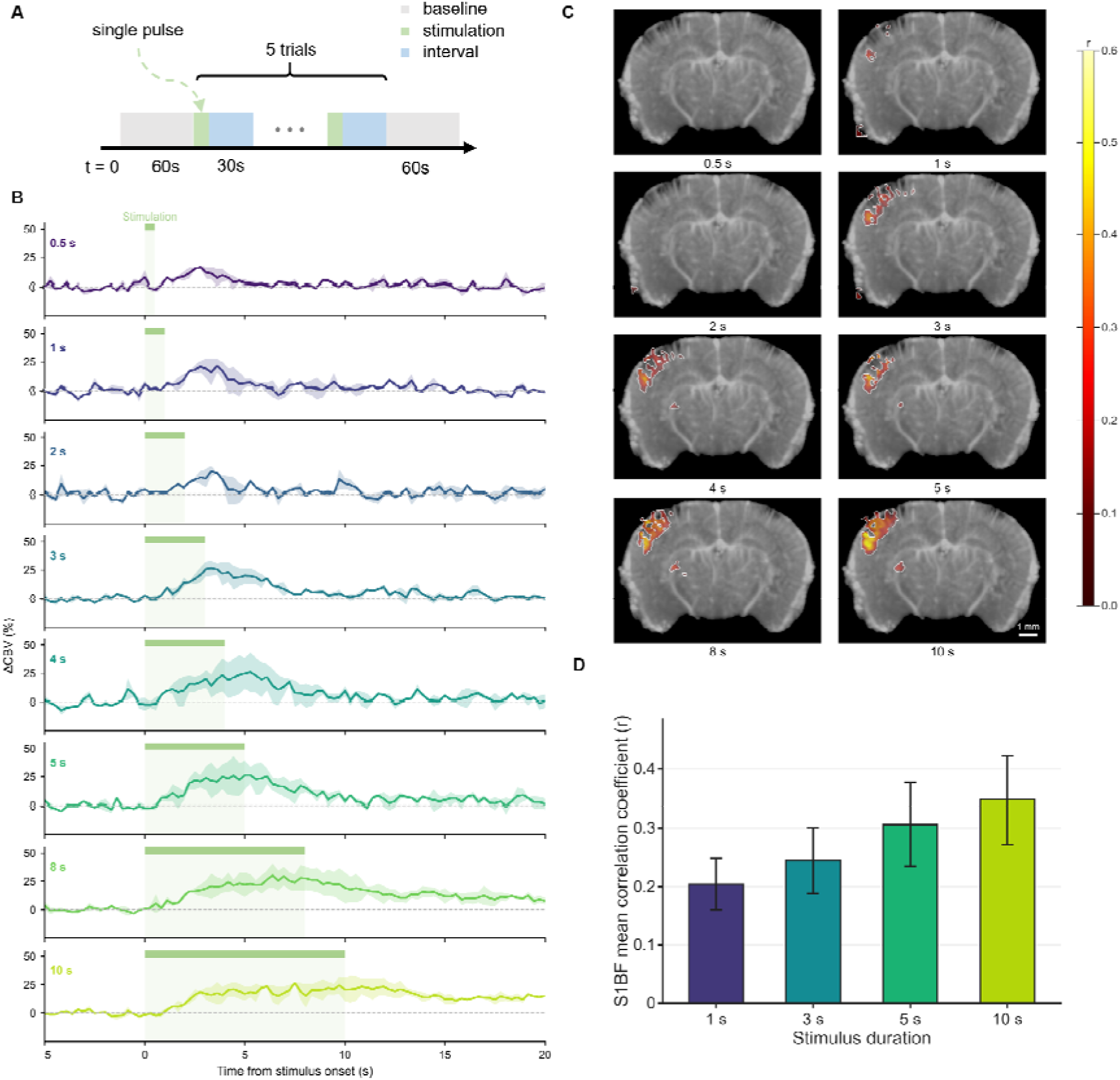
Duration-related characteristics of whisker-evoked fUS-CBV responses. **(A)** Schematic of the single-pulse whisker-stimulation paradigm. Each condition comprised five trials following a 60-s baseline, with 30-s intervals between trials. **(B)** Stimulus-aligned S1BF ΔCBV responses across stimulation durations from 0.5 to 10 s. Lines and shaded regions indicate the mean and SEM, respectively; green bars and pale-green shading indicate the stimulation periods, mean ± SEM across animals (n = 2 mice). **(C)** Representative atlas-aligned activation maps for 0.5-, 1-, 2-, 3-, 4-, 5-, 8-, and 10-s stimulation. Color indicates the correlation coefficient (r) between the fUS-CBV signal and the duration-matched stimulation regressor. **(D)** Mean correlation coefficient within the active S1BF area of the representative 1-, 3-, 5-, and 10-s maps shown in C. Bars indicate the spatial mean, and error bars indicate the spatial SD across active S1BF pixels. Scale bars, 1 mm.

Representative atlas-aligned activation maps were compared for all eight stimulation durations of 0.5, 1, 2, 3, 4, 5, 8, and 10 s at the same coronal level (Fig. 3C). The maps showed duration-associated differences in the spatial extent and correlation strength of the contralateral cortical response. Thresholded activation was limited in the shortest-duration representative maps, whereas broader and higher-correlation patterns were present under several longer-duration conditions. These spatial differences were summarized by calculating the mean correlation coefficient across active pixels within the S1BF region for the representative 1-, 3-, 5-, and 10-s maps (Fig. 3D). The mean coefficient increased across the four displayed conditions, consistent with the corresponding differences in activation strength and spatial extent. The bars and error bars describe the spatial mean and spatial variability of active-pixel correlation coefficients within each representative map.

Together, the duration-series experiments show that whisker-stimulation duration was associated with systematic differences in the magnitude, persistence, and spatial distribution of the fUS-CBV response. Brief isolated stimuli produced measurable S1BF signals, whereas longer stimulation generally produced stronger and more sustained responses. The temporal traces, atlas-aligned correlation maps, regional correlation summaries provide complementary descriptions of how stimulation-duration information is expressed in the measured fUS-CBV signal. These findings characterize the detectability of brief sensory inputs and the broad duration-related response features retained in fUS-CBV measurements.

### 4. Separability of successive fUS-CBV responses across inter-stimulus intervals

Paired-pulse experiments examined whether fUS-CBV responses evoked by two successive 2-s whisker stimuli could be identified as distinct temporal components. The unstimulated interval between the two pulses was varied across 1, 2, 3, 5, and 8 s (Fig. 4A).

**Figure 4.**
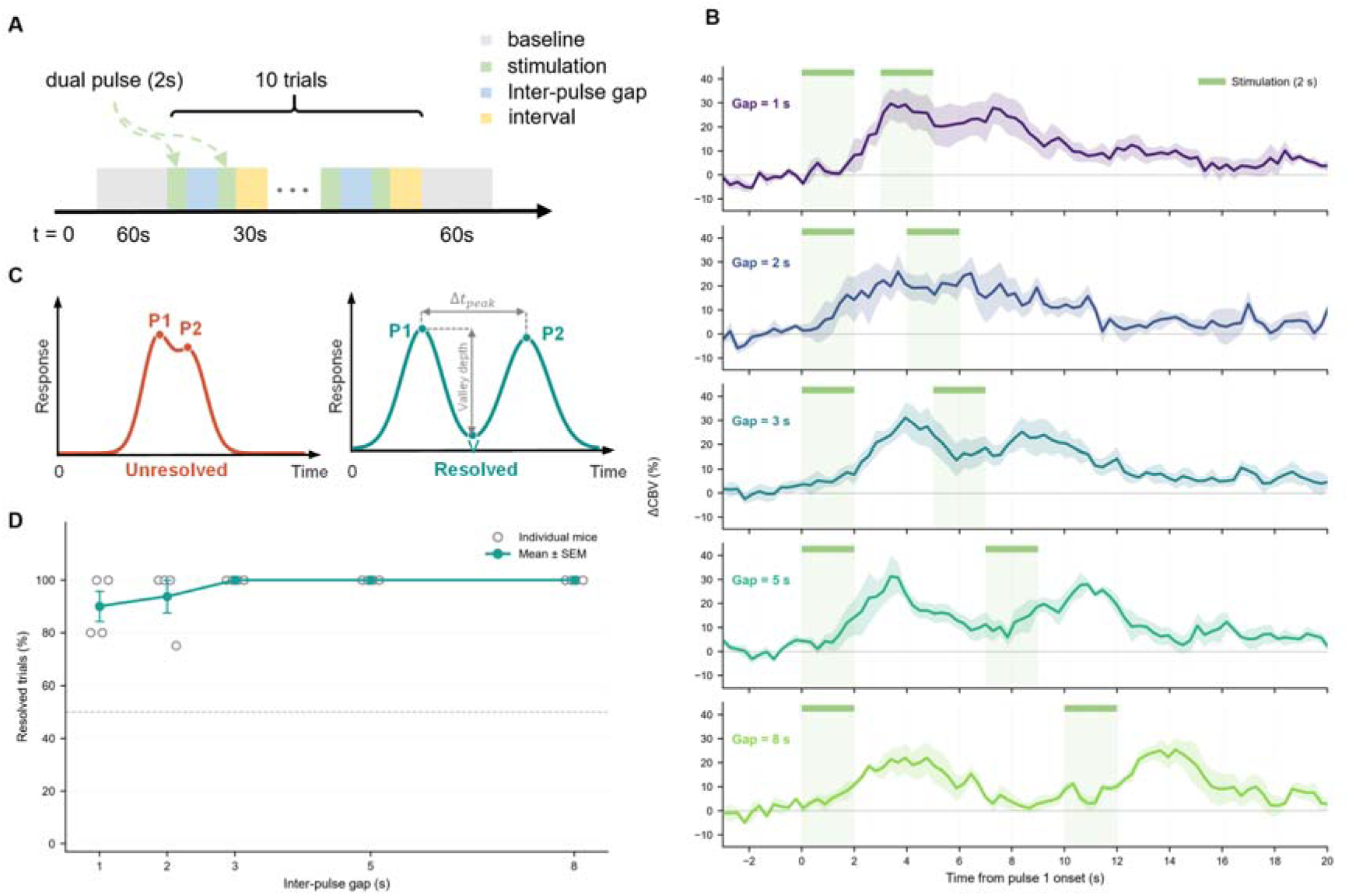
Temporal separability of successive whisker-evoked fUS-CBV responses. **(A)** Schematic of the paired-pulse paradigm. Two 2-s all-whisker stimuli were delivered with inter-stimulus intervals of 1, 2, 3, 5, or 8 s. **(B)** S1BF ΔCBV responses aligned to the onset of the first stimulus. Solid lines and shaded regions represent the mean and SEM across mice, respectively; green bars and shaded intervals indicate stimulation periods, mean ± SEM across mice (n = 4) **(C)** Schematic illustration of the response-separation criterion. Closely spaced peaks with an insufficient intervening valley were classified as unresolved, whereas two distinct peaks satisfying the predefined peak-separation and valley-depth criteria were classified as resolved. The curves are illustrative and do not represent experimental data. **(D)** Percentage of complete trials classified as resolved at each inter-stimulus interval. Gray open circles represent individual mice, and the colored line and symbols indicate the group mean ± SEM (n = 4 mice). The horizontal dashed line marks 50%. Detailed classification criteria are provided in Methods.

Stimulus-aligned S1BF ΔCBV traces from four mice showed two stimulus-associated response components whose temporal separation became more apparent as the inter-stimulus interval increased (Fig. 4B). At the shorter gaps, the intervening decline was less pronounced in the group-average waveform, whereas two response maxima were more clearly separated at the longer gaps. Group averaging can reduce trial-specific peak–valley structure when response timing varies across trials; paired-event separability was therefore evaluated from individual trials rather than from the group-average waveform alone.

Individual trials were classified using a predefined two-peak criterion (Fig. 4C). A trial was considered resolved when two local maxima were detected within the expected post-stimulus windows, the maxima were separated by at least 1 s, and the intervening valley showed sufficient absolute and proportional decline relative to the lower peak. The numerical thresholds used for classification are described in Methods.

Resolved-trial proportions were calculated separately for each mouse and summarized across four mice (Fig. 4D). Paired-event separability was already high at the shortest tested 1-s gap, increased further at a 2-s gap, and reached 100% at gaps of 3, 5, and 8 s.

Together, these results show that responses associated with successive 2-s whisker stimuli remained identifiable across the tested temporal spacings. Trial-level separability was high at the shortest 1-s inter-stimulus interval, and reached ceiling performance at gaps of 3 s or longer. These measurements provide characterization of paired-event separability and temporal conditions that can inform the decoding of successive sensory events from fUS-CBV signals.

### 5. Integrated spatial and temporal response characteristics relevant to fUS-CBV decoding

The preceding experiments characterized complementary spatial and temporal features of whisker-evoked fUS-CBV signals. Unilateral whisker stimulation produced contralaterally dominant responses in the S1BF and VPM. Stimulation of different individual whiskers generated distinguishable atlas-aligned activation distributions, while stimulation involving a greater number of whiskers was associated with broader cortical recruitment. These results indicate that whisker identity and stimulation extent were represented in the spatial organization of the measured fUS-CBV signals.

The duration-series and paired-pulse experiments characterized two aspects of temporal response expression. Isolated stimuli ranging from 0.5 to 10 s produced measurable group-averaged S1BF responses, with stimulus-duration-related differences in response magnitude, persistence and spatial correlation strength. For successive stimuli, paired-event separability was already high at the shortest tested 1-s inter-stimulus interval and reached 100% at gaps of 3 s or longer. The representation of an isolated stimulus and the separation of responses to successive stimuli therefore provide complementary information about the temporal characteristics retained in fUS-CBV signals.

Figure 5 qualitatively integrates the spatial and temporal properties examined in the preceding experiments. The horizontal dimension represents the degree to which whisker-associated cortical response distributions remain distinguishable, while the vertical dimension represents the separability of responses to successive stimuli across the tested temporal spacings. These dimensions describe complementary forms of stimulus-related information: spatial patterns can retain information about whisker identity and stimulation extent, whereas temporal response structure can retain information about isolated stimulus duration and the occurrence of successive events. The resulting organization summarizes the combinations of spatial and temporal information accessible in fUS-CBV measurements.

**Figure 5.**
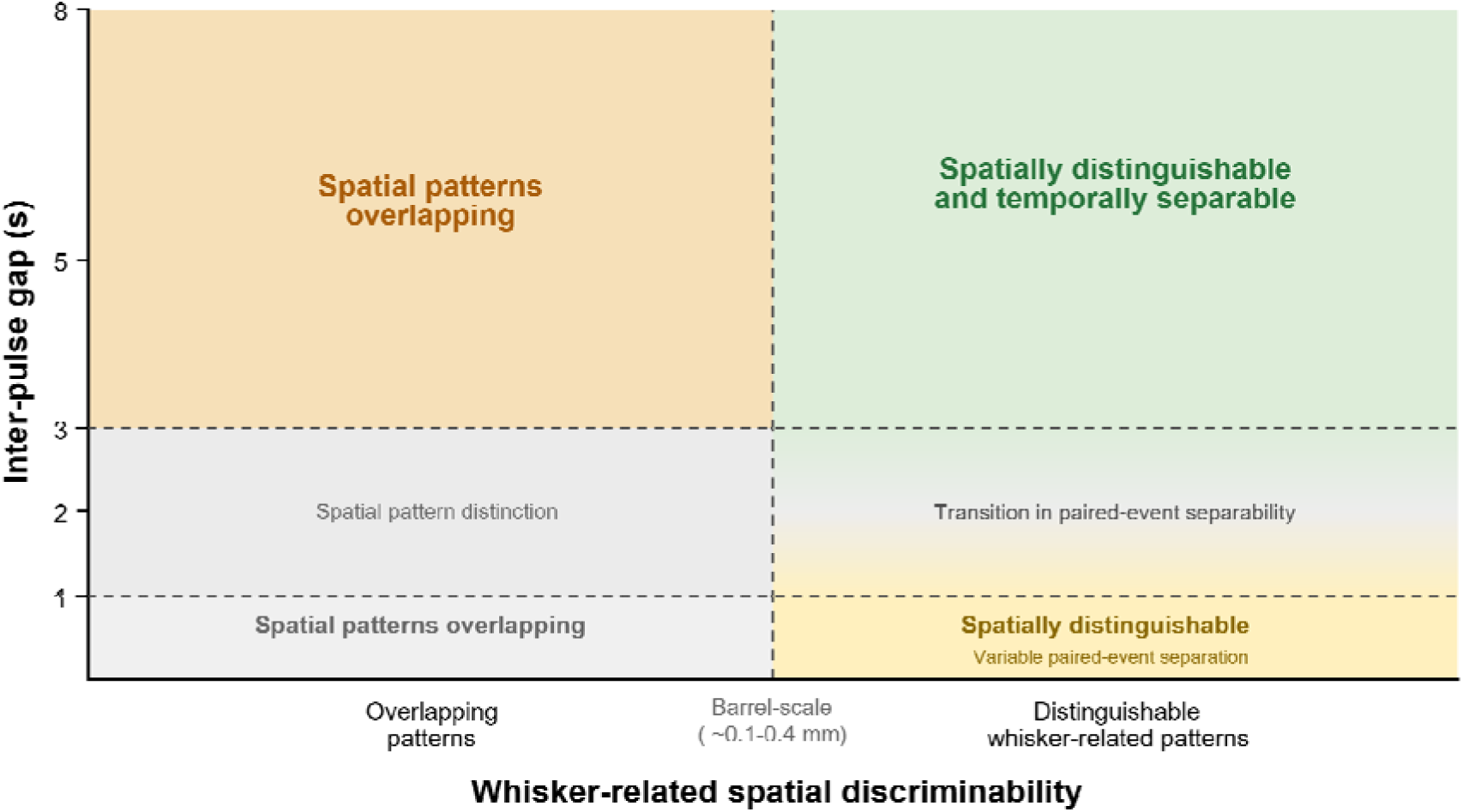
Integration of spatial pattern discriminability and temporal response characteristics in whisker-evoked fUS-CBV signals. The horizontal dimension represents the distinction among whisker-associated cortical response distributions. The vertical dimension summarizes the temporal information examined through isolated stimuli of different durations and paired stimuli presented at different inter-stimulus intervals. The four regions organize combinations of spatial pattern distinction and paired-event separability within the tested stimulation paradigms. The indicated mouse barrel diameter provides anatomical context based on published histological measurements of the mouse barrel cortex^29^.

Together, these findings show that decoding-relevant information in fUS-CBV signals is expressed through several complementary spatial and temporal response properties. Their experimental characterization provides a basis for selecting stimulation conditions, interpreting measured response patterns, and defining the spatial and temporal constraints used in the encoding and decoding analyses presented below.

### 6. Experimental response characteristics inform a proof-of-concept fUS-CBV encoding and decoding framework

The spatial and temporal response characteristics identified in the preceding experiments were organized into a proof-of-concept, experimentally informed framework linking controlled whisker-stimulation factors to measured fUS-CBV features (Fig. 6A). The encoding direction estimates spatial activation patterns and paired-response separability from whisker condition and inter-stimulus interval, whereas the decoding direction estimates stimulus-duration class from the measured S1BF ΔCBV time course with known stimulus onset. The framework therefore connects three stimulus factors—whisker condition, stimulus duration and inter-stimulus interval—with three corresponding signal features: spatial activation distribution, temporal response profile and paired-event separability.

**Figure 6.**
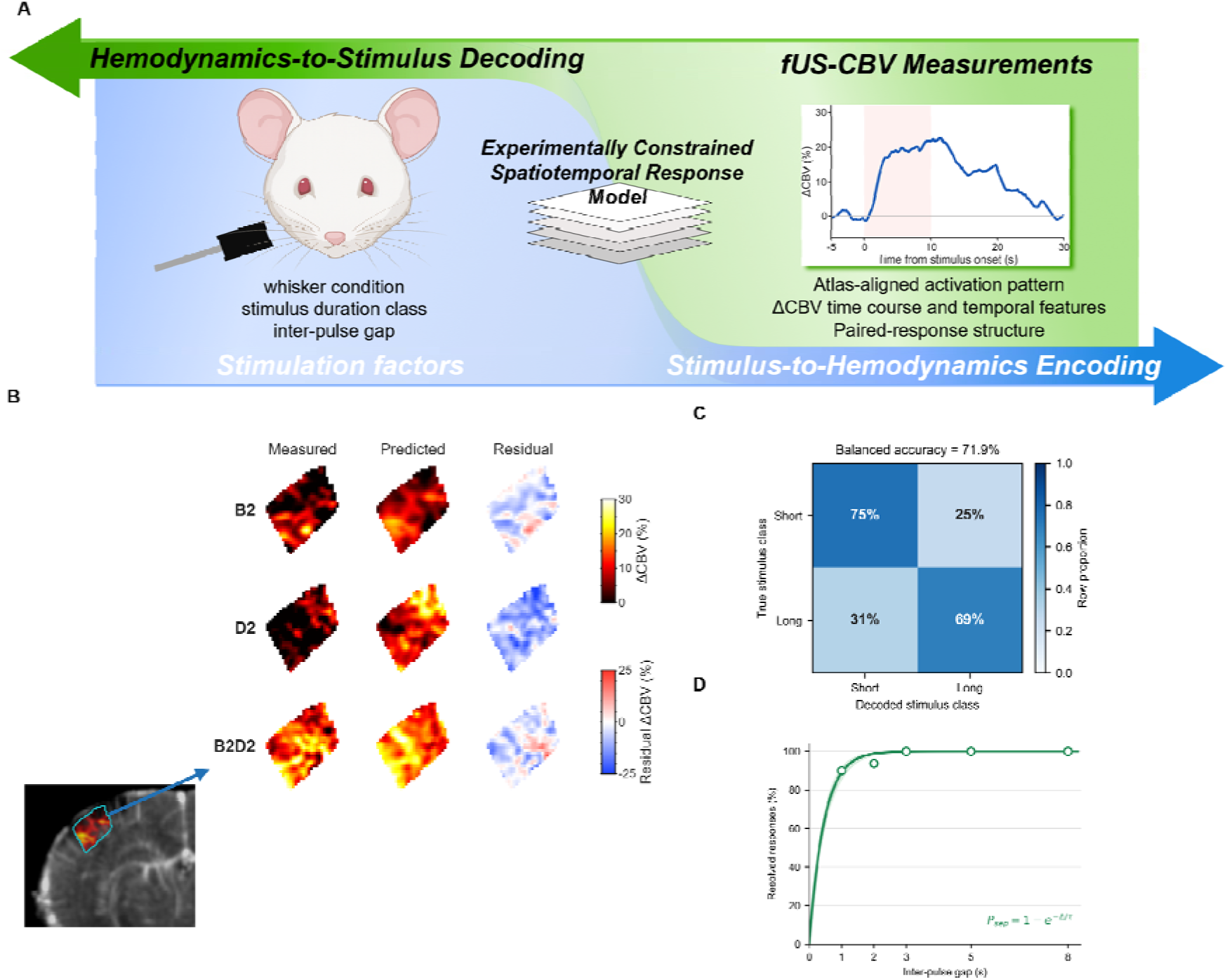
Experimentally constrained bidirectional framework for fUS-CBV encoding and stimulus decoding. **(A)** Conceptual framework linking whisker condition, stimulus-duration class, and inter-stimulus interval with spatial and temporal fUS-CBV features. The encoding direction predicts selected fUS-CBV response features from stimulus factors, whereas the decoding direction estimates selected stimulus-related properties from measured responses. **(B)** Reciprocal cross-animal prediction of atlas-aligned S1BF ΔCBV maps for B2, D2 and combined B2+D2 stimulation using the PCA-ridge spatial model. Columns show measured maps, predicted maps and residuals calculated as measured minus predicted ΔCBV. **(C)** Trial-level classification of short (≤2 s) and long (≥3 s) stimulation using fixed-HRF template matching. The row-normalized confusion matrix summarizes 229 retained trials from two mice; the balanced accuracy was 71.9%. **(D)** Relationship between inter-stimulus interval and paired-event separability. Open circles show the animal-equal mean resolved-trial proportions across four mice, the solid curve shows the fitted single-parameter recovery function as defined in Methods, and the shaded region indicates the range of leave-one-animal-out fits.

The spatial module evaluated reciprocal cross-animal prediction of condition-related activation patterns (Fig. 6B). Atlas-aligned trial-level ΔCBV maps from one animal were used to estimate low-dimensional spatial patterns associated with B2, D2 and combined B2+D2 stimulation, and the fitted model was then applied to the corresponding conditions in the other animal. The training and test animals were subsequently exchanged. Figure 6B presents the measured and predicted condition-related spatial distributions together with residual maps showing the remaining differences in response magnitude and spatial organization.

The temporal decoding module classified trial-level S1BF ΔCBV responses into short-duration (≤2 s) and long-duration (≥3 s) stimulus classes using time courses aligned to the known stimulus onset (Fig. 6C). The confusion matrix showed that 75% of short-stimulus trials and 69% of long-stimulus trials were assigned to the corresponding duration class, yielding a balanced accuracy of 71.9%. These results show that the temporal profiles of the measured fUS-CBV signals retained information associated with the broad duration category of the whisker stimulus.

The paired-event module summarized the relationship between inter-stimulus interval and the probability of identifying two response components (Fig. 6D). The resolved-trial proportion was already high at the shortest tested 1-s gap and reached 100% at gaps of 3, 5, and 8 s. A single-parameter phenomenological function summarized the measured relationship between inter-stimulus interval and paired-event separability.

Together, the three modules connect the experimentally characterized spatial and temporal response properties with selected encoding and decoding operations. The framework captures condition-related spatial patterns, broad stimulus-duration information and the recovery of paired-event separability using parameters derived from the corresponding experimental measurements. It provides a compact demonstration of how experimentally measured fUS-CBV response properties can inform defined stimulus-related prediction tasks.

## Discussion

This study characterized how spatial and temporal features of controlled whisker stimulation were represented in fUS-CBV signals. The experiments addressed three properties relevant to stimulus decoding: the distinction among whisker-associated spatial response patterns, the detectability and duration-related characteristics of isolated brief stimuli, and the separability of responses to successive stimuli. These properties describe which features of sensory input remained accessible after transformation through neurovascular coupling and vascular dynamics. The findings also establish experimental constraints that can guide the design of sensory-stimulation paradigms and the development of fUS-based encoding and decoding approaches.

Unilateral all-whisker stimulation produced contralaterally dominant fUS-CBV responses in the S1BF and VPM, establishing that the experimental platform captured stimulus-associated signals in cortical and thalamic somatosensory regions. Under individual-whisker stimulation, the atlas-aligned response distributions associated with C2 and D2 occupied neighboring but distinguishable cortical locations. Conditions involving more stimulated whiskers also showed broader response distributions in the representative maps. These observations are consistent with the ordered population-level organization of whisker representations in the barrel cortex ^21,22,26,29^. The spatial patterns measured by fUS reflect the combined effects of recruited neural populations, vascular architecture, and neurovascular coupling ^4–7^. Whisker identity was consequently expressed through distributed cortical response patterns with partial spatial overlap. Atlas registration and serial coronal mapping provided a common anatomical reference for comparing these patterns across imaging planes and experiments.

The duration-series experiments showed that isolated whisker stimuli ranging from 0.5 to 10 s produced measurable S1BF fUS-CBV responses. Response magnitude, persistence, and spatial distribution varied across stimulus durations, with longer stimulation generally producing stronger and more sustained signals. The response following 0.5-s stimulation establishes the detectability of a brief isolated sensory input under the present paradigm. Event detectability represents one aspect of temporal information, while the distinction between temporally adjacent events is addressed separately by the paired-stimulation experiments. The prolonged fUS-CBV response relative to the mechanical input is consistent with the temporal transformation of neural activity through neurovascular coupling and vascular response dynamics. The duration-series results therefore show that fUS-CBV time courses retain information about broad differences in stimulus duration and provide an experimental basis for classifying short- and long-duration stimulation.

The paired-stimulation experiments further characterize the temporal organization of fUS-CBV signals. When two 2-s stimuli were separated by a 1-s inter-stimulus interval, the corresponding response components frequently appeared as closely spaced or partially merged peaks. The proportion of trials satisfying the predefined two-peak criterion increased at a 2-s gap, and consistently high separability was observed at gaps of 3 s and longer across the four animals. These findings identify a transition from variable to reliable paired-event separation over the tested 1–3 s interval. The result is relevant to experimental designs in which successive sensory events are represented by overlapping CBV signals. Inter-pulse spacing influences whether individual events remain identifiable in the measured time course and therefore affects event labeling, response attribution and the interpretation of rapidly repeated stimulation paradigms.

The spatial and temporal experiments together show that the functional information accessible in fUS-CBV signals has several distinct dimensions. Spatial discriminability describes whether different whisker inputs retain distinguishable activation distributions. Brief-stimulus response detectability describes whether an isolated short input produces a measurable signal and whether broad duration-related differences remain accessible. Paired-event separability describes whether two temporally adjacent inputs produce identifiable response components. Figure 5 integrates the spatial and paired-event dimensions and summarizes the experimentally observed operating ranges of the present paradigm. A strength of this study is that these properties were examined through controlled changes in stimulation parameters using a consistent acquisition and analysis framework. The combination of atlas-aligned activation maps, ROI time courses, trial-level event classification and animal-level comparisons provides complementary evidence for interpreting spatial and temporal information in fUS-CBV data.

The experimentally informed encoding and decoding framework in Figure 6 translates these observations into a compact computational form. Whisker-associated spatial patterns provide the basis for cross-animal prediction of stimulus-specific activation maps. Duration-related response characteristics support classification of short- and long-duration stimulation from fUS-CBV time courses. The relationship between inter-stimulus interval and paired-event separability provides a recovery function for estimating whether two responses are likely to remain distinguishable. These modules connect experimentally measured properties with defined prediction tasks while retaining a limited number of interpretable parameters. This framework may support future inference of stimulus identity, duration class and temporal pattern from fUS-CBV measurements, as well as forward prediction of stimulus-associated spatial and temporal responses. Such capabilities could contribute to fUS-based brain–computer interfaces, optimization of sensory-stimulation paradigms and the development of data-informed functional brain models.

The present experiments cover selected whisker conditions, stimulus durations and inter-pulse intervals within a fixed imaging preparation. Extending the dataset to additional animals, matched coronal planes, intermediate stimulus durations and broader whisker combinations will allow the observed relationships to be estimated with greater coverage. Incorporating volumetric and longitudinal fUS measurements may further characterize how stimulus information is distributed across cortical and subcortical regions. Natural whisking and active-exploration paradigms could also test whether the experimentally derived spatial and temporal features remain informative under less periodic sensory conditions. Larger independent datasets will support further evaluation of cross-animal model transfer and refinement of the encoding and decoding framework.

In summary, controlled variation of whisker identity, whisker number, stimulus duration and inter-pulse interval revealed complementary spatial and temporal characteristics in fUS-CBV signals, characterizing the fundamental boundaries of fUS to reliably distinguish the neural activities information. Individual-whisker stimulation produced distinguishable activation distributions, brief isolated stimuli generated measurable responses with duration-related features, and paired-event separability increased as the inter-stimulus interval was extended. These findings provide systematic reference ranges for designing and interpreting whisker-evoked fUS experiments. Their integration into a compact computational framework also provides a basis for stimulus-state decoding, forward response prediction and future fUS-informed functional brain models.

## Conclusion

In conclusion, this study systematically characterizes how whisker identity, the number of stimulated whiskers, stimulus duration and inter-stimulus interval are represented in fUS-CBV signals. The results reveal distinguishable spatial distributions associated with individual-whisker stimulation, measurable responses to brief stimuli with duration-related characteristics, and progressively improved separability of paired responses as the inter-stimulus interval increases. Integrating these response characteristics into an experimentally informed encoding and decoding framework further demonstrates the feasibility of inferring stimulus states from fUS-CBV signals and predicting their corresponding hemodynamic responses. Collectively, these findings systematically characterize the spatiotemporal performance boundaries within which neural activity can be distinguished by fUS, providing an empirical basis and prior constraints for the design of future experimental paradigms and decoding models.

## Methods

### 1. Animals and experimental design

Male C57BL/6J mice aged 8–10 weeks were obtained from Bestest Biotechnology (Zhuhai, China). Animals were maintained under a standard light–dark cycle with free access to food and water. All procedures were approved by the Committee for the Use of Laboratory Animals of the Guangdong Institute of Intelligence Science and Technology and were conducted in accordance with institutional guidelines.

Cranial-window implantation was performed under isoflurane anesthesia using aseptic procedures. After removal of the scalp and exposure of the skull, a craniotomy was made over the target imaging region while preserving the dura mater. The removed bone was replaced with a 0.125-mm-thick polymethylpentene film (TPX), which was sealed at the margins and secured with dental cement to provide a stable acoustic window. A custom head-fixation mount was implanted when required for probe positioning and repeated imaging. Cefotaxime sodium (5 μg g^−1^) was administered intraperitoneally for three consecutive days after surgery. Animals were allowed to recover for at least 7 days before imaging. All fUS recordings were subsequently performed in awake, head-fixed mice without anesthesia.

The experimental design examined how controlled variations in whisker stimulation were represented in fUS-CBV signals. Unilateral all-whisker block stimulation was first used to characterize lateralized responses in cortical and thalamic somatosensory regions. Subsequent experiments varied whisker identity and the number of stimulated whiskers to examine spatial response distributions. Isolated stimuli ranging from 0.5 to 10 s were used to characterize duration- related response features, while paired-pulse experiments examined the separability of responses to two successive 2-s stimuli across different inter-stimulus intervals. These experiments provided the spatial and temporal measurements used in the encoding and decoding framework.

Some animals contributed to more than one imaging session or stimulation paradigm. For fixed-plane experiments, each dataset was analyzed within its predefined target coronal plane. Serial coronal-plane acquisitions were retained as separate spatial measurements for atlas-aligned mapping. Repeated trials were treated as within-animal observations, and the animal was used as the biological unit for group-level statistical comparisons.

### 2. Functional ultrasound imaging and stimulus synchronization

Functional ultrasound data were acquired using a custom mini-fUS system^30^ equipped with a 15-MHz, 64-element linear-array transducer with an element pitch of 0.1 mm. The transducer was positioned above the TPX cranial window and oriented in the coronal plane using a fixed probe holder. Medical-grade ultrasound coupling gel was applied between the transducer and the cranial window to maintain acoustic coupling throughout each recording.

Power Doppler image sequences were reconstructed from ultrafast ultrasound acquisitions using plane-wave compounding and clutter filtering. Power Doppler intensity was used as a relative measure of local cerebral blood volume, consistent with established fUS imaging principles. The reconstructed power Doppler data were sampled at 3.6 Hz and stored as three-dimensional matrices with dimensions of 181 × 127 × T, corresponding to depth, lateral position, and time. The reconstructed image-grid spacing was approximately 0.05 mm, providing a lateral field of view of approximately 6.4 mm.

For serial spatial mapping, consecutive coronal planes were acquired at approximately 0.1-mm intervals along the anterior–posterior axis. Each plane was retained as an individual spatial measurement and subsequently assigned to its corresponding atlas position. Experiments examining stimulus-duration-related responses and paired-event separability were conducted at fixed coronal planes containing the target somatosensory region.

Whisker stimulation was controlled by an Arduino-based system and temporally synchronized with fUS acquisition. A synchronized trigger signal was recorded for each acquisition and used to align the stimulation sequence with the power Doppler time series. Each recording included a pre-stimulation baseline, one or more stimulation periods, and a poststimulation recovery period, with the exact timing determined by the corresponding stimulation paradigm described below.

### 3. Whisker stimulation paradigms

Mechanical whisker stimulation was delivered to the right whiskers using an Arduino-controlled servo motor coupled to a custom soft-contact applicator. The applicator consisted of a compliant contact element mounted on a rigid support arm and was positioned to engage the selected whisker or whisker group without contacting the face. During stimulation, the commanded servo position alternated between 0° and 45° at 4 Hz, producing repeated whisker deflections. The stimulation frequency and servo excursion were maintained across experiments, while whisker identity, the number of stimulated whiskers, stimulus duration, and inter-stimulus interval were varied according to the experimental paradigm.

For the initial response-characterization and spatial-mapping experiments, stimulation was applied to selected individual whiskers, a predefined two-whisker combination, or the intact whisker array. The conditions used in the main spatial analyses included C2 and D2 stimulation for comparing individual-whisker response distributions, as well as B2, combined B2+D2, and intact all-whisker stimulation for comparing representative cortical response distributions across different stimulation extents. Each acquisition began with a 60-s baseline period, followed by five 20-s stimulation blocks separated by 40-s unstimulated intervals. Serial coronal-plane acquisitions were used to examine the spatial distribution of the stimulus-associated fUS-CBV signals.

For the stimulus-duration experiments, the intact whisker array was stimulated for 0.5, 1, 2, 3, 4, 5, 8, or 10 s. Stimulation frequency and servo excursion were unchanged across duration conditions. Each recording began with a 60-s baseline period, and each duration condition comprised five repeated trials separated by 30-s unstimulated recovery intervals. Duration-series measurements were acquired at fixed coronal planes containing the contralateral S1BF. The same target plane was maintained across the duration conditions acquired within a given imaging series.

For the paired-pulse experiments, each trial consisted of two 2-s stimulation trains delivered to the intact whisker array. The inter-stimulus interval was defined as the interval between the end of the first stimulation train and the onset of the second and was set to 1, 2, 3, 5, or 8 s. Each gap condition was repeated within each animal to obtain trial-level S1BF fUS-CBV responses. Four mice contributed to the paired-pulse analysis, and the numbers of complete trials retained at each inter-stimulus interval are reported in the Results and the corresponding figure legend.

### 4. fUS signal processing and activation mapping

Power Doppler image sequences were processed using custom MATLAB scripts. Signals outside the imaged brain region were excluded using anatomical brain masks. Relative cerebral blood volume changes were calculated from the power Doppler intensity as

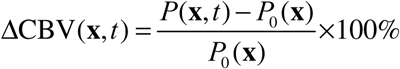

where *P*(**x**,*t*) is the power Doppler intensity at spatial location **x** and time *t*, and *P*_0_(**x**) is the mean power Doppler intensity during the corresponding pre-stimulation baseline period. For repeated-trial experiments, baseline normalization was performed using the pre-stimulation period associated with each trial before trial-level responses were combined.

Voxel-wise activation maps were generated by calculating the Pearson correlation coefficient between each fUS-CBV time series and a stimulus-related response regressor. The regressor was constructed from the recorded stimulation sequence and convolved with a predefined response kernel to account for the delayed temporal profile of the fUS-CBV signal. For experiments with different stimulation durations, the onset and duration of the corresponding stimulus condition were incorporated into the regressor. The voxel-wise correlation coefficient was calculated as

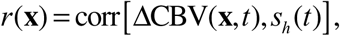

where *r*(**x**) is the correlation coefficient at location **x**, and *t* is the temporally modeled stimulus regressor. Positive correlation values within the brain mask were retained for activation mapping. A common correlation scale and activation criterion were applied when maps from different conditions were compared directly. For the comparison of single B2, combined B2+D2, and intact all-whisker stimulation in Fig. 2C, activated pixels were defined using a common threshold of *r* ≥ 0.15.

Anatomical registration was performed using the vascular power Doppler image from each coronal plane as the native-space reference. The vascular image was aligned to the corresponding coronal section of the Allen Mouse Brain Common Coordinate Framework ^31^ using anatomical landmarks visible in the fUS image and atlas reference. The resulting spatial transformation was applied without further adjustment to the corresponding correlation maps and ΔCBV maps. Atlas-aligned images were assigned to their anterior–posterior coordinates relative to Bregma. For serial spatial datasets, consecutive coronal sections were assembled according to their atlas positions to generate two-dimensional montages and three-dimensional representations of the activation distribution.

For comparisons between individual whisker conditions, atlas-aligned activation maps were displayed at matched coronal positions using common spatial coordinates and correlation scales. Dual-color overlays were generated by assigning the D2- and C2-associated correlation maps to separate color channels, allowing their spatial distributions to be visualized within the same atlas-aligned field.

### 5. Spatiotemporal response analyses

Anatomically guided regions of interest were defined for the contralateral and ipsilateral primary somatosensory barrel field (S1BF) and ventral posteromedial thalamic nucleus (VPM) using the atlas-aligned anatomy and the corresponding fUS vascular image. Mean power Doppler signals were extracted across all pixels within each ROI and converted to ΔCBV time courses using the corresponding pre-stimulation baseline. Repeated trials acquired at the same target coronal plane were aligned to stimulus onset and averaged within each animal. For group-level time-course summaries, each animal contributed one animal-level trace, and the group mean and SEM were calculated across animals.

For the hemispheric comparisons shown in Fig. 1E, the correlation coefficient between the ROI fUS-CBV signal and the modeled stimulation regressor was calculated separately for the contralateral and ipsilateral S1BF and VPM. When more than one predefined eligible measurement was available from an animal, the correlation coefficients were Fisher-z transformed and summarized to obtain one value per ROI and animal. Contralateral and ipsilateral animal-level values were compared using two-sided paired t-tests. The S1BF and VPM comparisons were corrected for multiple testing using the Holm method.

Spatial response distributions associated with whisker identity were evaluated from atlas-aligned correlation maps. D2- and C2-associated maps were displayed within the same atlas coordinate system using separate color channels to compare their cortical locations. Representative maps from single B2, combined B2+D2, and intact all-whisker stimulation were compared at selected atlas-aligned coronal planes. A common correlation scale and activation threshold of r ≥ 0.15 were applied to the three displayed conditions.

For the stimulus-duration analysis, retained trials from one predefined target coronal plane per animal were aligned to stimulus onset and averaged separately for each duration condition. Animal-level traces were then combined with equal weighting to obtain the mean S1BF ΔCBV response and SEM shown in Fig. 3B. Correlation maps for the all eight duration conditions of 0.5, 1, 2, 3, 4, 5, 8, and 10 s conditions were evaluated at the same atlas-aligned coronal level and displayed in Fig. 3C. For the representative 1-, 3-, 5-, and 10-s maps summarized in Fig. 3D, the mean correlation coefficient was calculated across active pixels within the atlas-defined S1BF region. Error bars in Fig. 3D indicate the spatial SD across these pixels within each representative map.

Paired-event separability was assessed from individual S1BF ΔCBV trials using a predefined two-peak procedure. The first and second response peaks were searched within 1.5–6.0 s after the onset of their corresponding stimuli, and the intervening minimum was identified between the two peaks. Candidate peaks were required to be separated by at least 1 s. A strict separation criterion required an intervening valley-to-lower-peak ratio of no more than 0.735, an absolute valley depth of at least 0.5 ΔCBV percentage points, and a relative valley depth of at least 10% of the lower peak above baseline. Peak prominence, the decline from the first peak to the valley, and the rebound from the valley to the second peak were additionally evaluated to identify traces with a consistent two-response morphology. The classification procedure was applied independently to the original trace and a trace smoothed using a 1-s moving-average window. A trial was classified as resolved when the original and smoothed traces both supported two distinguishable response components. Trials with insufficient peak separation, valley depth, or response rebound were classified as unresolved. Borderline trials and trials with inconsistent classifications between the original and smoothed traces were retained in the denominator but were not counted as resolved. The resolved-trial proportion was calculated separately for each animal and inter-stimulus interval. Group values in Fig. 4D are presented as the mean ± SEM of the animal-level resolved-trial proportions.

Trials with incomplete acquisition windows or prominent signal artifacts identified by the predefined quality-control procedure were excluded before response averaging or trial classification. No individual time points were removed from otherwise retained trials.

Unless otherwise specified, group data are presented as mean ± SEM across animals. Statistical analysis and figure generation were conducted using MATLAB 2024b and GraphPad Prism 10.4.1.

### 6. Experimentally Informed fUS-CBV Encoding and Decoding Framework

An experimentally informed, proof-of-concept framework was constructed to describe selected relationships between whisker-stimulation factors and fUS-CBV response features. The framework contained three components: cross-animal prediction of spatial ΔCBV patterns, decoding of stimulus-duration class from S1BF ΔCBV time courses, and estimation of paired-response separability from the inter-stimulus interval. Each component was fitted using the corresponding experimental dataset and generated the outputs presented in Fig. 6.

For the spatial component, retained trial-level ΔCBV maps from B2, D2, and combined B2+D2 stimulation were transformed into the same atlas-aligned coronal plane. Each spatial ΔCBV map was calculated by averaging ΔCBV over 5–30 s after stimulus onset and was vectorized within the common S1BF analysis mask. The resulting matrix was denoted by **M**∈□*^n×p^*, where *n* is the number of training trials and *p* is the number of spatial locations retained within the common mask. For one training animal, the resulting maps formed a matrix **M**, in which each row represented one trial and each column represented one spatial location. The centered map matrix was defined as

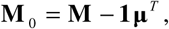

where **μ** is the mean spatial map of the training animal. Here, **1** ∈ □*^n^* is a column vector of ones, **μ** ∈ □*^p^* is the mean spatial map represented as a column vector, and superscript *T* denotes matrix transpose. Principal component analysis was applied to the centered maps, and the retained spatial components formed the basis matrix **V***_K_*. The corresponding trial scores were

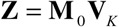

Here, *K* is the number of retained spatial components, **V***_K_* ∈ □*^p×K^* contains the corresponding spatial basis vectors, and **Z** ∈ □*^n×K^* contains the PCA scores of the training trials.

The relationship between stimulation condition and spatial scores was estimated using ridge regression:

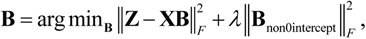

The design matrix **X**∈□^*n*×4^ contained an intercept, an indicator for B2 involvement, an indicator for D2 involvement, and an interaction indicator for simultaneous B2+D2 stimulation. The corresponding coefficient matrix was denoted by **B**∈□^4×*K*^. *λ* controls the regularization applied to the non-intercept coefficients.

For stimulation condition *c*, the predicted activation map was reconstructed as

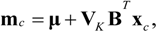

where **x***_c_* represents the corresponding stimulation condition. The model was fitted using data from one animal and applied to the other animal, after which the training and test animals were exchanged. The number of retained spatial components and the regularization strength were selected using trial-wise validation within the training animal. Data from the test animal were excluded from model fitting and parameter selection. Condition-label permutation was used to assess whether the observed cross-animal spatial correspondence exceeded that obtained after disrupting the relationship between stimulation condition and activation pattern.

For stimulus-duration decoding, the input was the retained trial-level S1BF ΔCBV time course *y_i_*(*t*), evaluated over 0–30 s after the known stimulus onset. Candidate response templates were generated by convolving a boxcar stimulus of duration *d* with a fixed single-gamma hemodynamic response kernel:

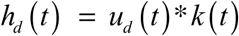

where *u_d_*(*t*) is the stimulus function and *k*(*t*) is the fixed hemodynamic kernel. A non-negative gain was fitted separately for each trial and candidate duration:

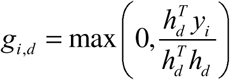

The estimated duration was the candidate producing the lowest normalized root-mean-square error:

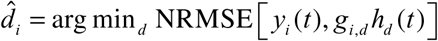

The estimated durations were grouped into short (≤ 2 s) and long (≥ 3 s) classes. Classification was summarized using a row-normalized confusion matrix and balanced accuracy. This analysis therefore estimated stimulus-duration class conditional on a known stimulation onset.

For paired-event separability, each trial was classified as resolved or unresolved according to the predefined two-response criterion described above. The resolved-trial proportion was calculated separately for each animal and inter-stimulus interval. The relationship between inter-stimulus interval Δ and the probability of resolving two responses was represented by

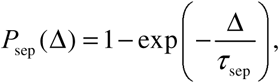

where *P*_sep_ (Δ) is the estimated separability probability and *τ*_sep_ is the fitted recovery time constant. The parameter was estimated by minimizing the mean squared error across the animal-level observations, with each animal contributing equally at each tested gap. The fitting was repeated after excluding each animal in turn to assess the stability of the estimated separability curve.

## Acknowledgments

This work was supported in part by the National Key Research and Development Program of Ministry of Science and Technology of China (2023YFC2410900),National Natural Science Foundation of China (32371151), Hong Kong Research Grants Council Collaborative Research Fund (C5053-22 GF), General Research (15126524 and 15224323), internal funding from the Hong Kong Polytechnic University (G-SACD), Research Center for Non-invasive Brain Computer Interface (1-CE0M), and Research Institute of Smart Ageing (1-CDJM).

## Data Availability

The processed source data generated in this study are available in the Zenodo repository at https://doi.org/10.5281/zenodo.22283977.

## Code Availability

Custom MATLAB and Python code used for data processing, figure generation, and the encoding and decoding analyses is available at https://github.com/bcbc1998/Whisker_fUS_reproducibility_code_v2. The repository includes documentation and scripts for reproducing the source-data visualizations for Figures 1–6 from the processed source-data package.

## Author Contributions

C.B. and J.Y. contributed to study design, experiments, data analysis, software development, visualization, and manuscript writing and revision. M.X. contributed to experiments and data analysis. Y.H.S.S. contributed to data analysis, visualization, and manuscript writing and revision. J.H., J.Z., and Z.C. contributed to data processing, validation and manuscript revision. H.-Y. L, Y.W, Z.Y. and Z.Q. provided resources, funding, supervision, and manuscript revision. All authors reviewed and approved the final manuscript.

## Competing Interests

The authors declare no competing interests.

## Notes

### Competing Interest Statement

The authors have declared no competing interest.

### Summary of Updates

This revised version includes updated analyses and figures that clarify the spatial response patterns, duration-related signal characteristics, and paired-event separability of whisker-evoked fUS-CBV signals. An experimentally informed proof-of-concept encoding and decoding framework has been added. The Methods, figure legends, references, and manuscript formatting have also been revised to improve methodological clarity, consistency, and reproducibility. Processed source data and reproducibility code are now available through Zenodo and GitHub.

